# Network cloning using DNA barcodes

**DOI:** 10.1101/098343

**Authors:** Sergey A. Shuvaev, Batuhan Başerdem, Anthony Zador, Alexei A. Koulakov

## Abstract

The connections between neurons determine the computations performed by both artificial and biological neural networks. Recently, we have proposed SYNseq, a method for converting the connectivity of a biological network into a form that can exploit the tremendous efficiencies of high-throughput DNA sequencing. In SYNseq, each neuron is tagged with a random sequence of DNA—a “barcode”—and synapses are represented as barcode pairs. SYNseq addresses the analysis problem, reducing a network into a suspension of barcode pairs. Here we formulate a novel and complementary synthesis problem: How can the suspension of barcode pairs be used to “clone” or copy the network back into an uninitialized tabula rasa network? Although this synthesis problem might be expected to be computationally intractable, we find that, surprisingly, this problem can be solved efficiently, using only neuron-local information. We present the “one barcode one cell” (OBOC) algorithm, which forces all barcodes of a given sequence to coalesce into the same neuron, and show that it converges in a number of steps that is a power law of the network size. Rapid and reliable network cloning with single synapse precision is thus theoretically possible.

---

The connections between neurons determine the computations performed by a neural network. In both biological and artificial neural networks, connections are established and tuned by experience and learning. Connections can thus be considered a “summary” of the statistical structure of the experience—data—on which the network was trained. This summary may be considerably more compact and efficient than the original data. For example, deep neural networks for object recognition contain tens of millions of connections derived from training sets consisting of hundreds of billions pixels, which results in more than 1000-fold compression [1, 2]. It would therefore be more efficient to copy these connections onto a new network than to retrain a new network from scratch.

Most current implementations of artificial neural networks exploit digital computers and GPUs [2]. On these architectures, connections are stored explicitly and are therefore straightforward to extract and copy into a new network. In biological networks, by contrast, there is no central repository for connections, so reading out the connections of a network and copying them into a new network represents a difficult challenge. During neural development, for example, a genomic DNA sequence representing prior evolutionary experience is converted into the brain’s connectivity. Similar challenges may arise in future artificial or hybrid biological/artificial architectures.

We have recently proposed SYNSeq, a novel approach to determining neuronal connectivity [3, 4]. The key idea is to convert the connections into a form that can be read out using high-throughput DNA sequencing, thereby benefitting from the advances in sequencing technology. Sequencing is now extremely fast and inexpensive—it is routine to decode billions of DNA fragments per day, and sequencing cost has dropped at a rate faster than Moore’s law. To convert neuronal connectivity into a sequencing problem, we induce individual neurons to express unique random nucleotide identifiers called “barcodes.” Pairs of pre- and postsynaptic barcodes represent individual synaptic connections. These barcode pairs can then be used to represent the connectivity of a network (Fig. 1).

**Figure 1.**
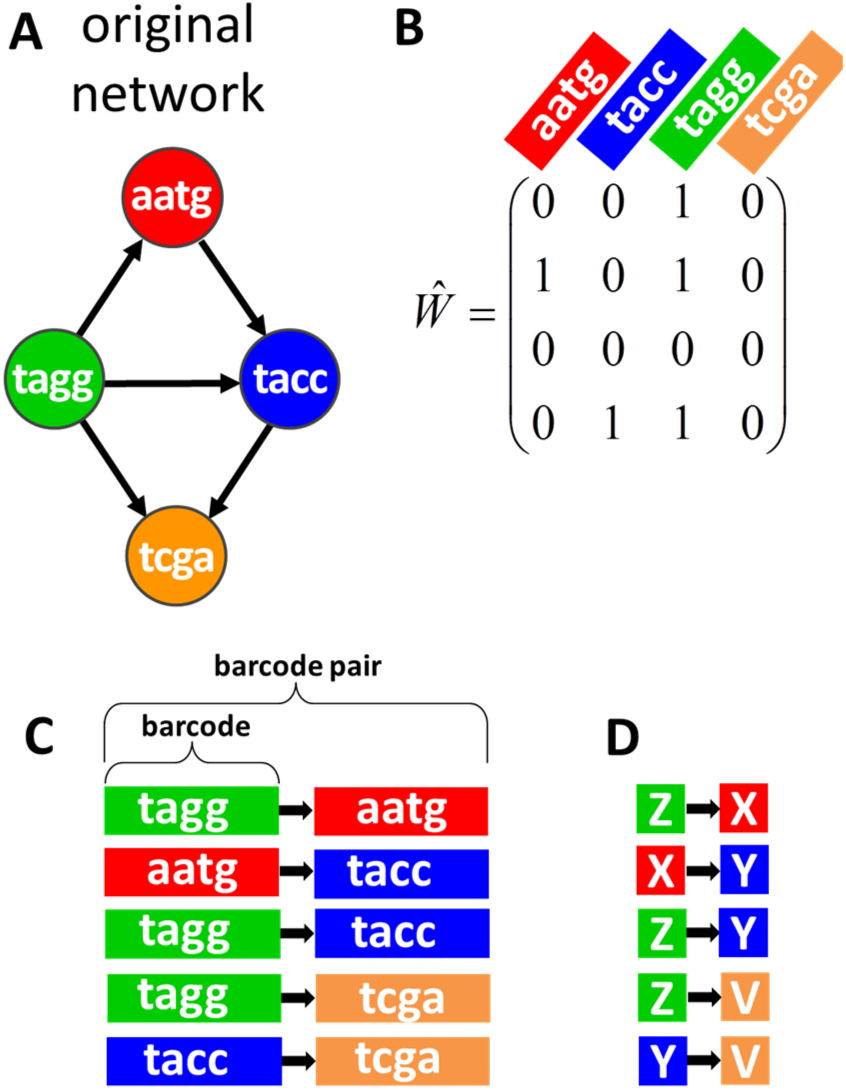
Representation of a network by an ensemble of barcode pairs (SYNSeq). (A) An example of small network. In SYNSeq, each neuron is represented by a short unique nucleotide sequence called barcode. (B) The connectivity matrix corresponding to the network in (A). (C) Network connections are encoded by pairs of barcodes with a spacer (black arrow) representing the connections’ direction. (D) We represent barcodes by unique letters of an alphabet for brevity.

Here we formulate a different problem: Given an ensemble of connections represented by barcode pairs, can we copy them into a new network? In other words, can the original network be cloned? We explore a computational model that simulates the behavior of barcodes introduced into a *tabula rasa* network with unstructured connectivity and test its ability to recreate target connectivity in such networks. We require that the underlying mechanisms be purely local, i.e. that the algorithm uses only information available to a given neuron and its synapses. Below we present an algorithm that allows robust copying of connectivity based only on local interactions.

In our approach, connectivity is specified by unique molecular labels (DNA barcodes) with single-synapse precision. It is commonly assumed that implementing connectivity via individual synaptic tags is not feasible due to the absence of guidance mechanism that would direct barcodes to the right synapses [5]. One might expect that establishing desired connectivity using individual synaptic labels would require a number of steps that is exponential in network size. The inadequacy of unique molecular tags in instructing connectivity had motivated Roger Sperry to introduce the idea of molecular gradients [6]. Here, we find, surprisingly, that a form of molecular dynamics proposed by us yields convergence to the target connectivity in polynomial in network size number of steps, even though the connectivity is specified by unique molecular labels for each synapse. This finding implies that copying connectivity with single-neuron precision using our strategy is theoretically possible.

Our algorithm attempts to recreate the target connectivity between neurons (Fig. 1A). The connectivity can be represented as a connection matrix *Ŵ* (Fig. 1B). We assume that every network node (neuron) is identified by a unique barcode, *i.e.* by a sequence of nucleotides long enough to label uniquely every neuron in the network (Fig. 1A). Network connectivity is thus encoded by barcode pairs, where each barcode pair consists of a presynaptic barcode, a postsynaptic barcode, and a spacer between them indicating the connection’s direction (Fig. 1C). The number of barcode pairs is equal to the number of non-zero entries in the connection matrix, or to the total number of connections in the network. To simplify notation, we will represent each barcode by a single letter of the alphabet rather than as a string of nucleotides (Fig. 1D).

The barcode pairs are introduced into synapses of a *tabula rasa* network that is, initially, fully connected (Fig. 2B). Since connectivity in our model is directional, we assume that, between every two cells, synapses are formed initially in both directions. The full connectivity assumption is made here to simplify the description of network dynamics, and is reminiscent of the overproduction of synaptic connections that occurs during development [7, 8] and the full potential cortical connectivity [9]. The number of neurons in the *tabula rasa* network is assumed to be equal to the number of nodes in the desired network, i.e. equal to the number of barcodes.

**Figure 2.**
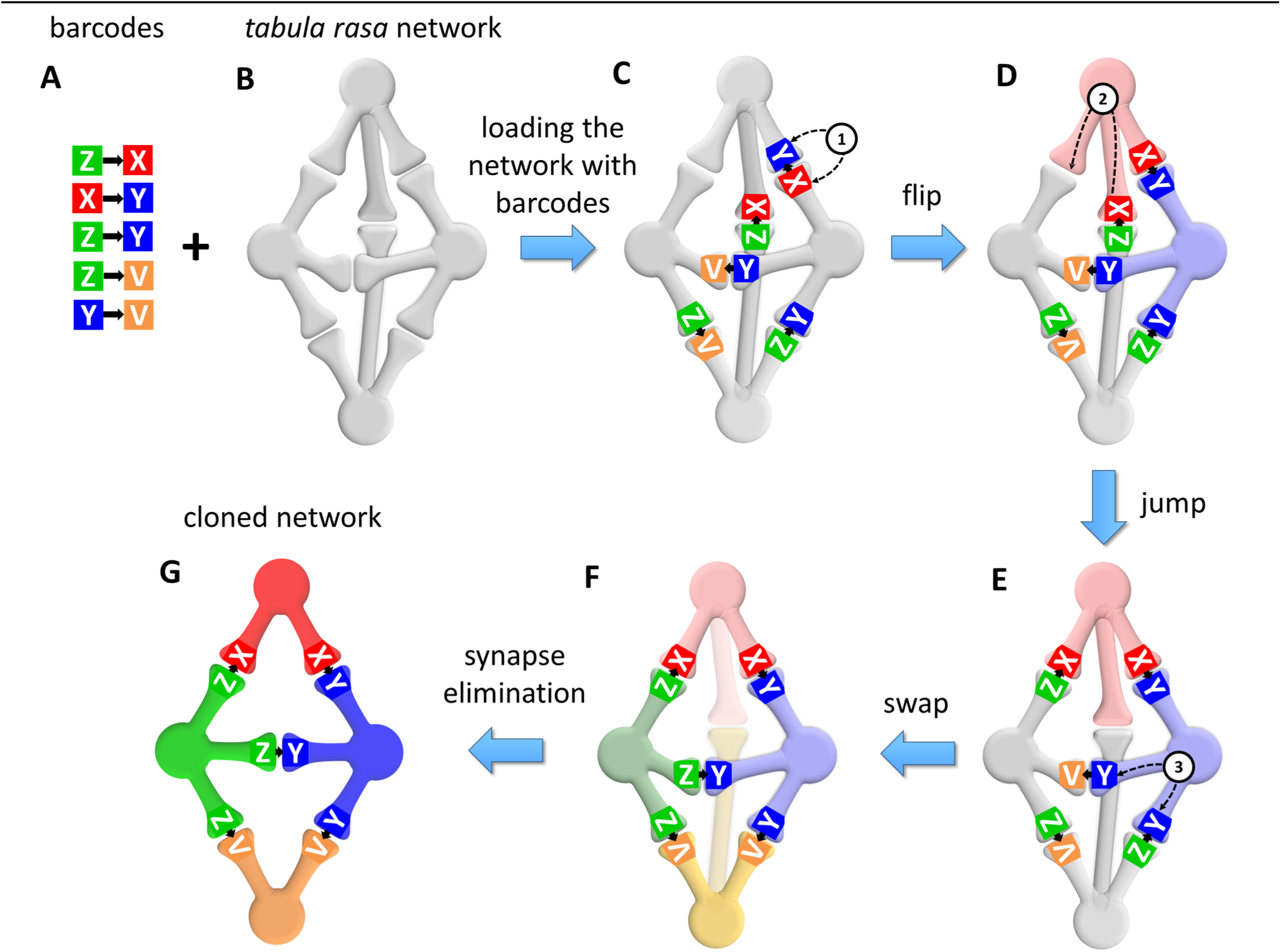
The “one barcode one cell” (OBOC) rule yields target connectivity. (A) The set of barcode pairs representing the original network from Fig. 1. Individual barcode sequences are shown as letters for brevity. Barcode pairs represent individual synapses. (B) An all-to-all connected *tabula rasa* network that receives the ensemble of barcode pairs. We show connections as undirected synapses for simplicity. (C) The barcode pairs are initially arranged randomly. (D-F) Barcode pairs can move through the network by jumping from synapse to synapse using three moves as illustrated: jumps (1), flips (2), and swaps (3). The moves minimize the cost function defined by equation (1). (F) Minimization of the cost-function forces all barcodes facing every neuron to be the same. This arrangement is called OBOC. Once OBOC solution is achieved, we eliminate all synapses that contain no barcode pairs, such as the synapse between cells “X” and “V”.(G) OBOC solution yields the copying of the original connection matrix.

The barcodes are initially introduced into synapses randomly. The barcodes are then rearranged in the network via three types of moves. First, each barcode can be reinserted in the synapse between the same pair of cells in different orientation (“flips”; Fig. 2C). Second, the barcodes can jump from one synapse to another synapse of the same cell (“jumps”; Fig. 2D). Finally, two barcodes located in the same neuron can trade places?

(“swaps”; Fig. 2E). To practically implement these three moves, we select two synapses of the same neuron at random, ensure that at least one of them contains a barcode pair, and swap the pairs, even if source and destination are the same or one of them is empty. In implementing these moves we keep track of the direction of barcode pairs and synapses, i.e. barcode pairs are introduced into synapses of the correct orientation. We ensure that the described moves are local in that the barcode pairs are only relocated between synapses of the same neuron.

Using this set of moves, we rearrange barcode pairs in the network attempting to implement the “one barcode – one cell” (OBOC) solution. In the OBOC solution all barcodes in the synapses of the same cell, facing this cell, are the same (Fig. 2G). Thus, in Fig. 2G, *all* barcodes in the rightmost cell are described by letter Y (V, X, Y, Z is a short-hand notation for much longer nucleotide sequences). Similarly, all barcodes in the leftmost cell are labeled by letter Z. We reasoned that if the logic of the interaction of cells and barcodes favors OBOC solution, cells will discover their identity as encoded by barcodes. Because every cell in the *tabula rasa* network has a potential to become any cell as defined by the barcodes, a specific arrangement of barcode pairs respecting OBOC rule is associated with a symmetry breaking, whereby the network selects one possible assignment of barcodes into cells out of *N* ! combinations *(N* is the number of neurons in the network, and is equal to the number of barcodes). We also reasoned that if we then eliminate all synapses that are *not* occupied by a barcode pair, the remaining synapses will implement the target connectivity.

To practically implement OBOC solution, we defined a cost-function, *H*, that is minimized by the barcode dynamics. The cost function depends on the synapse-barcode connection index (SBCI), *χ_ij,vμ_*, which determines which barcode pair is present in what synapse. This variable is equal to 1 or 0 if a barcode pair connecting two barcodes *μ* → *v* is present or absent in a synapse from cell *j* to *i* ( *μ*, *v*, *i* and *j* are unique indexes enumerating barcodes and cells). The constraint on SBCI is that after summing it over all synapses, we should obtain the original barcode connectivity matrix shown in Fig. 1B: 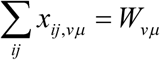. Index 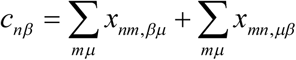
defines the number of barcodes (not pairs) of type *β* in cell number *n*. Although many choices are possible for the cost function, we use this particular form:

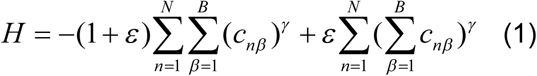

Here sums are assumed over the neuronal index *n* ranging from 1 to *N*, the total number of neurons, and the barcode index *β*, ranging from 1 to *B*, the total number of barcodes. *γ* and *ε* are the parameters of the cost-function. To implement the OBOC rule, the parameter γ must exceed unity. In the present work we used *γ* = 2, in which case the first term of the cost-function can be viewed as a measure of the sparseness of barcode distribution *c_nβ_* [10, 11]. Minimizing a measure determined by barcode sparseness will converge to a single barcode dominating each cell, i.e. to the desired OBOC solution. The second term in the cost-function defines the penalty for placing a nondominant barcode in each cell. This penalty is controlled by an independent parameter, *ε*, which we set to *ε* = 10 here. For *γ* = 2, the cost function can be written as 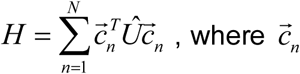 is the vector of barcode abundances in neuron number *n*, and *Û* = −(1 + *ε*)*Î* + *εŶ*. Here *Ŷ* is the matrix of all ones. Because the diagonal part of matrix *Û* (˜ *Î*) is negative, it favors solutions in which there is a single barcode type per cell, while the off-diagonal part (˜*Ŷ*) penalizes multiple barcode types in a cell. Both of these components help achieve OBOC solution. Importantly the cost function has a property of locality, i.e. the contribution for each neuron depends on the variables available to this neuron *c_nβ_*, i.e. the number of barcodes of type *β*.

The approach based on minimizing a cost-function is one of the ways to quantitatively describe biological processes and has been used successfully to describe establishing connectivity, especially when competition or interdependence between cells is important [12, 13]. To minimize the cost function we use Metropolis Monte Carlo (MMC) simulated annealing procedure [12, 13] and three types of barcode moves as described above. After the cost function is minimized, at the end of MMC procedure, we remove synapses that carry no barcodes. Overall, we hypothesized that, when OBOC solution is reached and empty synapses are eliminated, the final connectivity between cells will reproduce the connectivity between barcodes.

Our results show that, indeed, OBOC rule yields desired connectivity after several MMC steps (Fig. 3). We tested the convergence on a set of randomly generated asymmetric sparse networks, with the fraction of nonzero connections determined by parameter *f*. We find that, even for substantially large connection matrices (Fig. 3D-F), the target connectivity can be reached in relatively small number of steps in 100% of cases. To quantify the speed of convergence, for each MMC simulation, we computed the number of attempts to move barcodes before a perfect OBOC solution was achieved, *N*_steps_. We find that this parameter is well approximated by a power-law function

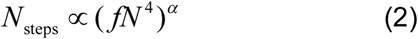

**Figure 3.**
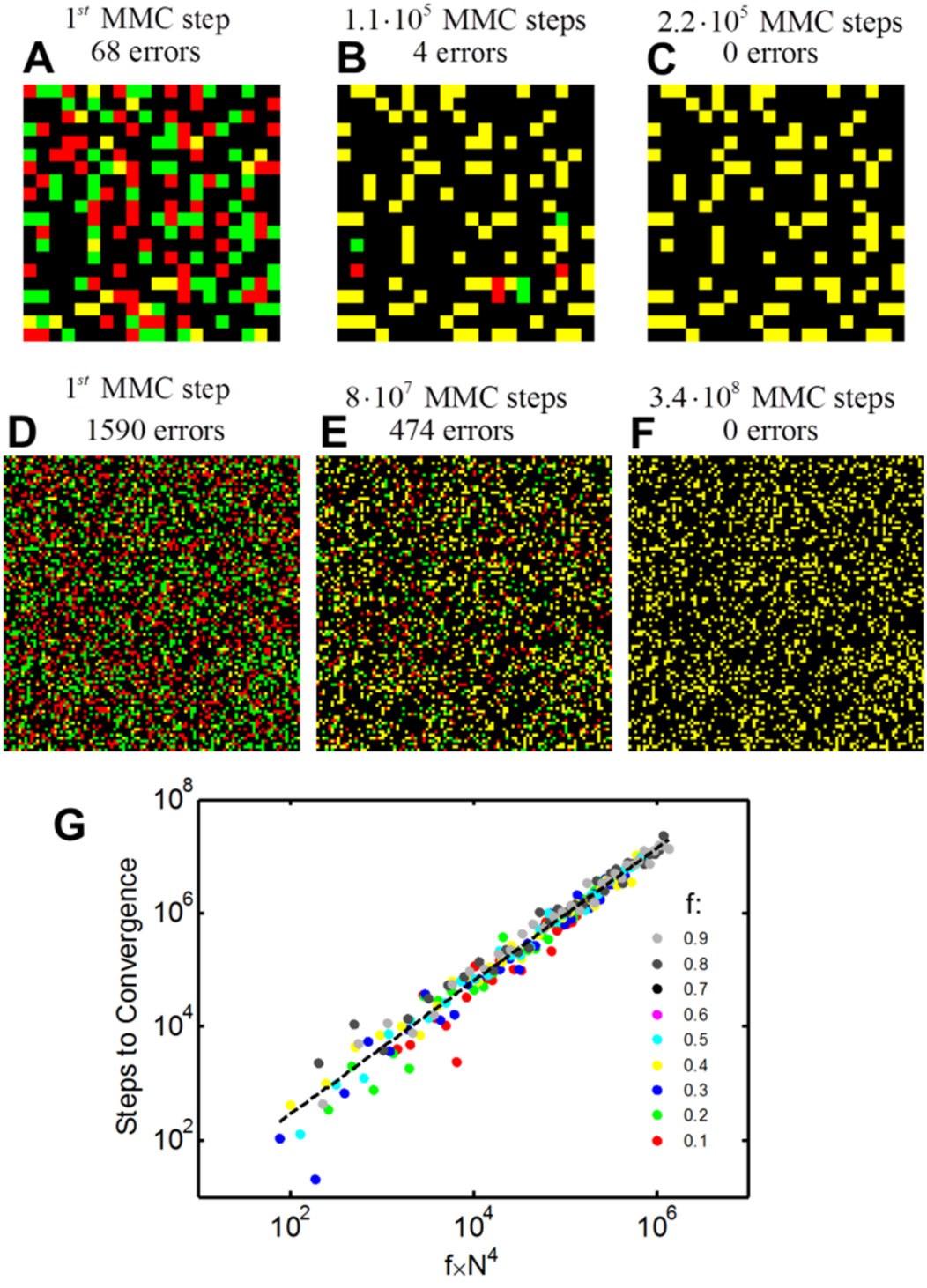
The OMOC rule allows copying desired network connectivity matrix. Results of a single MMC simulated annealing run for 20x20 (A-C) and 100x100 (D-F) networks. Red/green channels show target/actual connection matrices. Yellow matrices at the end of the simulation run (C and F) indicate a perfect copy. (G) OBOC rule yields target connectivity in a number of steps given by a power law of network size. Number of steps required for convergence as a function of a combination of network parameters (*N*, the size of the network, and *f*, the fraction of non-zero connections). Dashed line is the best linear fit corresponding to the number of steps Nsteps « *(fN*^4^)^117^

Here *N* is the number of cells in the network (the size of the connection matrix), *α* ∝ 1.17 is the scaling exponent. This finding means that OBOC rule yields target connectivity in ˜ *N*^4.68^ steps, i.e. a power law function of the network size. This result suggests that even for large networks, the original connectivity can be reached in a finite, i.e. non-exponential, number of steps. Overall our results show that copying the target connectivity to the new network is possible with OBOC rule, and that the convergence of this rule is relatively fast.

If one assumes that the scaling exponent *α* ∝ 1.17 is approximately equal to one, the number of steps to convergence can be represented as *N*_steps_ ≃ *BN*^2^, where *B* = *fN*^2^ is the total number of barcodes (synapses) in the network. The number of steps that each barcode has to make is given by *N*_steps_ / *B* ≃ *N*^2^. Thus, to a first approximation, each barcode has to explore ˜ *N*^2^ potential positions before OBOC solution is achieved, the number that is independent on other barcodes. This may explain why the rate of convergence is given by a power law function and is not exponential in the number of neurons.

Here we addressed the question whether connectivity can be copied from one neural network to another. We assumed that the connections are represented by an ensemble of DNA barcodes [3, 4]. We analyzed the dynamics of barcodes introduced into a clean slate *tabula rasa* network. The particular form of dynamics that we considered is described by one barcode one cell rule (OBOC), which favors positioning of a single type of barcodes in a single neuron. We showed that OBOC dynamics leads to fast and reliable recreation of desired connectivity in the new network. The formation of new connectivity is achieved in a number of steps given by a power law of the network size. Thus, copying connectivity from one neural network to another using DNA barcodes is theoretically possible.

## Methods

To minimize the cost function (1), we used simulated annealing [12, 13]. We started from a random distribution of barcodes in synapses of a fully connected directed network. Barcodes were relocated between synapses as described, according to Monte Carlo statistical rules. The temperature was gradually lowered from 10^−2^ to 10^−6^ of the initial value. The number of steps in the algorithm was chosen to be 10 times the value given by equation (2). The probabilities of three operations, jumps, swaps, and flips, were 1 – *f*, *f* −1/ *N*, and 1/ *N* respectively. During each of the operations, barcodes were inserted in a random orientation. To compare to the target connectivity, we used a greedy procedure that finds dominant barcodes for each cell.

